# Neural dynamics of causal inference in the macaque frontoparietal circuit

**DOI:** 10.1101/2021.12.06.469042

**Authors:** Guangyao Qi, Wen Fang, Shenghao Li, Junru Li, Liping Wang

**Affiliations:** Institute of Neuroscience, Key Laboratory of Primate Neurobiology, CAS Center for Excellence in Brain Science and Intelligence Technology, Chinese Academy of Sciences, Shanghai 200031, China; University of Chinese Academy of Sciences, Beijing 100049, China

## Abstract

Natural perception relies inherently on inferring causal structure in the environment. However, the neural mechanisms and functional circuits that are essential for representing and updating the hidden causal structure and corresponding sensory representations during multisensory processing are unknown. To address this, monkeys were trained to infer the probability of a potential common source from visual and proprioceptive signals on the basis of their spatial disparity in a virtual reality system. The proprioceptive drift reported by monkeys demonstrated that they combined historical information and current multisensory signals to estimate the hidden common source and subsequently updated both the causal structure and sensory representation. Single-unit recordings in premotor and parietal cortices revealed that neural activity in premotor cortex represents the core computation of causal inference, characterizing the estimation and update of the likelihood of integrating multiple sensory inputs at a trial-by-trial level. In response to signals from premotor cortex, neural activity in parietal cortex also represents the causal structure and further dynamically updates the sensory representation to maintain consistency with the causal inference structure. Thus, our results indicate how premotor cortex integrates historical information and sensory inputs to infer hidden variables and selectively updates sensory representations in parietal cortex to support behavior. This dynamic loop of frontal-parietal interactions in the causal inference framework may provide the neural mechanism to answer long-standing questions regarding how neural circuits represent hidden structures for body-awareness and agency.

## INTRODUCTION

The brain is constantly confronted with a myriad of sensory signals. Natural perception relies inherently on inferring the environment’s hidden causal structure(Deroy, Spence, & Noppeney, 2016; French & DeAngelis, 2020; Lochmann & Deneve, 2011). In the process of building representation of the bodily self, the brain combines, in a near-optimal manner, information from multiple sensory inputs. When a single entity (e.g. the bodily self) evokes correlated noisy signals, our brain combines the information to infer the properties of this entity on the basis of the quality and uncertainty of the sensory stimuli. As a result, behavioral performance often benefits from combining information using such uncertainty-based weighting across sensory systems(Stein & Stanford, 2008). However, in a natural environment, multiple sensory cues are typically produced by more than one source (for example, two entities), which should not be integrated in the brain, especially when the superposing cues are sufficiently dissimilar and uncorrelated. Instead, the brain’s inferential process of integration breaks down, leading to the perception that these cues originate from distinct entities. This process of inferring the causes of sensory inputs for perception is known as causal inference(Kording et al., 2007).

Thus far, most of neurobiological studies of multisensory processing have operated under the assumption that different streams of sensory information can arise from the same source. For example, previous neurophysiological research in monkeys showed that neurons implement reliability-weighted integration on the premise that visual and vestibular signals are from one common source(Fetsch, DeAngelis, & Angelaki, 2013; Morgan, Deangelis, & Angelaki, 2008; Porter, Metzger, & Groh, 2007). Therefore, despite the ubiquity of the phenomenon of causal inference and much psychophysical and theoretical research(Acerbi, Dokka, Angelaki, & Ma, 2018; Dokka, Park, Jansen, DeAngelis, & Angelaki, 2019; Kayser & Shams, 2015; Kording et al., 2007; Mohl, Pearson, & Groh, 2020; Rohe & Noppeney, 2015; Sato, Toyoizumi, & Aihara, 2007), its neural mechanisms and functional circuits remain largely unknown. While recent studies in humans have begun to establish neural correlates(Aller & Noppeney, 2019; Cao, Summerfield, Park, Giordano, & Kayser, 2019; Rohe, Ehlis, & Noppeney, 2019; Rohe & Noppeney, 2015, 2016), the sequential process of first encoding the sensory signals, subsequently combining them with prior information to infer whether the sources should be assigned to the same entity for later information integration or segregation, and finally, but most importantly, updating prior information for both the hidden structure of the environmental sources and their downstream sensory representations has never been studied at the single-neuron resolution in animals.

In the present study, we established an objective and quantitative signature of causal inference at a single-trial level using a reaching task and a virtual reality system in macaque monkeys. We showed that monkeys combined historical information and current multisensory signals to estimate the hidden common source, and more importantly, subsequently updated both the causal structure and sensory representation during the inference. We then further recorded from the premotor and parietal (area 5) cortices of three monkeys to investigate the neural dynamic and functional circuits of causal inference in multisensory processing. Our behavioral and neural results reveal a complete account of neural computation that appears to mediate causal inference behavior, which includes inferring a hidden common source and updating prior and sensory representations at different hierarchies.

## RESULTS

### Behavioral paradigm

Using a virtual-reality system, we trained three monkeys (monkeys H, N, and S) to reach for a visual target with their nonvisible (proprioceptive) arm while viewing a virtual arm moving in synchrony with a preset spatial visual-proprioceptive (VP) disparity (Fig. 1A). On each trial of the experiment, the monkeys were required to initiate the trial by placing their hand on the starting position (blue dot) for 1 s and were instructed not to move. After the initiation period, the starting point disappeared and the visual virtual arm was rotated; this mismatch arm was maintained for 0.5 s as the preparation period. The reaching target was presented as a “go” signal, and monkeys had to reach toward the visual target within 2.5 s and place their hand in the target area for 0.5 s, referred to as the target-holding period, to receive a reward. Any arm movement during the target-holding period automatically terminated the trial. The proprioceptive drift due to the disparity between visual and proprioceptive inputs was measured at the endpoint of the reach and was defined as the angle difference between the proprioceptive arm and the visual target (the estimated arm) (Fig. 1B, see details of animal training and reward in Methods). In addition to this VP-conflict (VPC) task, two control experiments were conducted: (i) where the visual and proprioceptive information were perfectly aligned (VP task) and (ii) where there was only a proprioceptive signal (P task). The procedures of the three tasks (VPC, VP, and P) were essentially identical, except that the visual or proprioceptive information presented to monkeys varied according to the context of the experiment (see Methods). Using a block design, the order of three different blocks (tasks) in each training or recording session was randomized.

**Figure 1.**
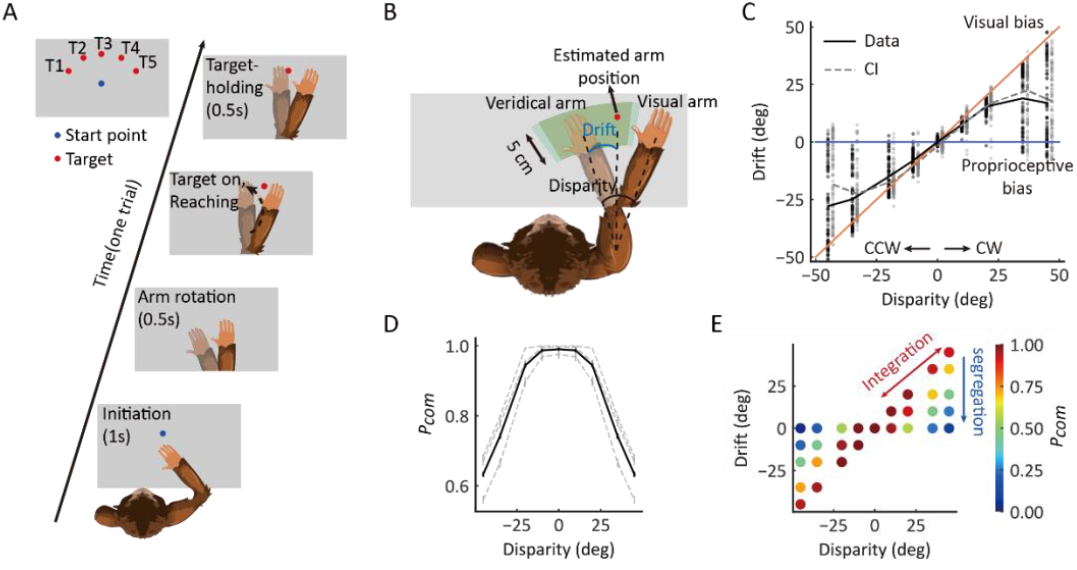
Behavioral task and proprioceptive drift results. **(A)** Overview of the behavioral task. The monkey was instructed to hold its proprioceptive arm over the starting position (blue dot) to initiate one trial. After the rotation of the virtual visual arm, a virtual red dot was presented, and the monkey was required to place its proprioceptive arm on the target and hold to get a reward. **(B)** Schematic drawing of reward area, proprioceptive drift, and the different types of arms (veridical/proprioceptive and virtual/visual). Here, proprioceptive drift was defined as the rotated degree from the veridical arm position to the estimated arm position (the same as the target location) measured from the shoulder. The reward area is defined by the green area, which ensured the monkey performed the task in a rational way and without visual feedback (see animal training in Methods). **(C)** Example behavioral results from one session of one monkey (also see Fig. S1). CCW, counterclockwise; CW, clockwise. The black line represents raw data. The gray line represents the BCI model fitting result. **(D)** The average *P*_*com*_ as a function of disparity. The black line represents the average *P*_*com*_ across monkeys. The dashed lines represent the average *P*_*com*_s across sessions of three monkeys separately. Error bars indicate standard errors of the means (SEMs). **(E)** Model prediction of the *P*_*com*_. Each point represents the average *P*_*com*_ in each cluster grouped by specific disparity and proprioceptive drift according to the monkey’s behavior.

### Causal inference framework and monkey’s behavior

The hierarchical Bayesian causal inference (BCI) model encodes probability distributions over the two sensory (visual and proprioceptive) signals and incorporates rules that govern how a prior belief about the sensory causal structure is combined with incoming information to judge the event probability in proprioception. To examine whether the monkeys inferred the causal structure during multisensory processing, we first examined the proprioceptive drift as a function of disparity in the VPC task. Overall, the three monkeys showed a very consistent behavioral pattern, with the proprioceptive drift increasing for small levels of disparity and plateauing or even decreasing when the disparity became larger (e.g., exceeded 20°) (Fig. 1C; for data on individual monkeys, see Fig. S1). The BCI model qualitatively explains the nonlinear dependence of drift as a function of disparity. For small disparities, there is a high probability that the proprioceptive and visual signals came from the same source. Hence, the visual information is fully integrated with the proprioceptive information. For large disparities, however, it is likely that the proprioceptive and visual signals are from different sources, leading to a breakdown of integration and consideration of only the proprioceptive information (segregation). In this case, visual information has a weak weight in the integration. As a consequence, the effect of disparity on drift is reduced. The BCI model quantified the nonlinear dependence between disparity and proprioceptive drift to measure the posterior probability of a common source (*P*_*com*_), the consequence of cause inference. We fitted the behavioral data using the BCI model. The results showed two signatures of the *P*_*com*_ pattern: (i) the averaged *P*_*com*_ decreased as the disparity increased (Fig. 1D), and (ii) within each disparity, especially the large ones, the *P*_*com*_ decreased as the proprioceptive drift decreased (Fig. 1E) (see individual monkeys’ behavior in Fig. S1).

More importantly, the model posits that not only the inference of the causal structure is based on visual and proprioceptive inputs but also the subsequent updating of (i) the prior belief of causal structure based on the historical information (e.g., probability of a common source in the previous trials), and (ii) the uncertainty of sensory signals for the visual and proprioceptive recalibration (Fig. 2A). To test these hypotheses, we first implemented the Markov analysis of the prior belief and *P*_*com*_ (see Methods) to see whether the prior probability of a common source (*P*_*prior*_) in the current trial depended on the historical *P*_*com*_ (Fig. 2B). The Markov model included the transition probability of *P*_*prior*_ between the current (n^th^) and previous (n^th^ − 1) trial to account for the trial-by-trial variability in spatial drifts observed in the three monkeys (Fig. 2B, left). The fit to the model demonstrated that the *P*_*com*_ observed in the n^th^ trial was significantly affected by that in the previous (n^th^ − 1) trial (Wilcoxon signed-rank test, *p* < 0.001), indicating that the *P*_*com*_ was computed on the basis of both *P*_*prior*_ from the previous trial and the sensory inputs, with their disparity, from the current trial. Note that the transition probabilities (*P*_*(C=1*|*C=1)*_ and *P*_*(C=1*|*C=2)*_) remained relatively high (larger than 0.8 in three monkeys) was because that overall the number of high *P*_*com*_ trial was much more than low *P*_*com*_ trial in either training or recording sessions. This was consistent with high baseline *P*_*prior*_ in three monkeys (Table S1).

**Figure 2.**
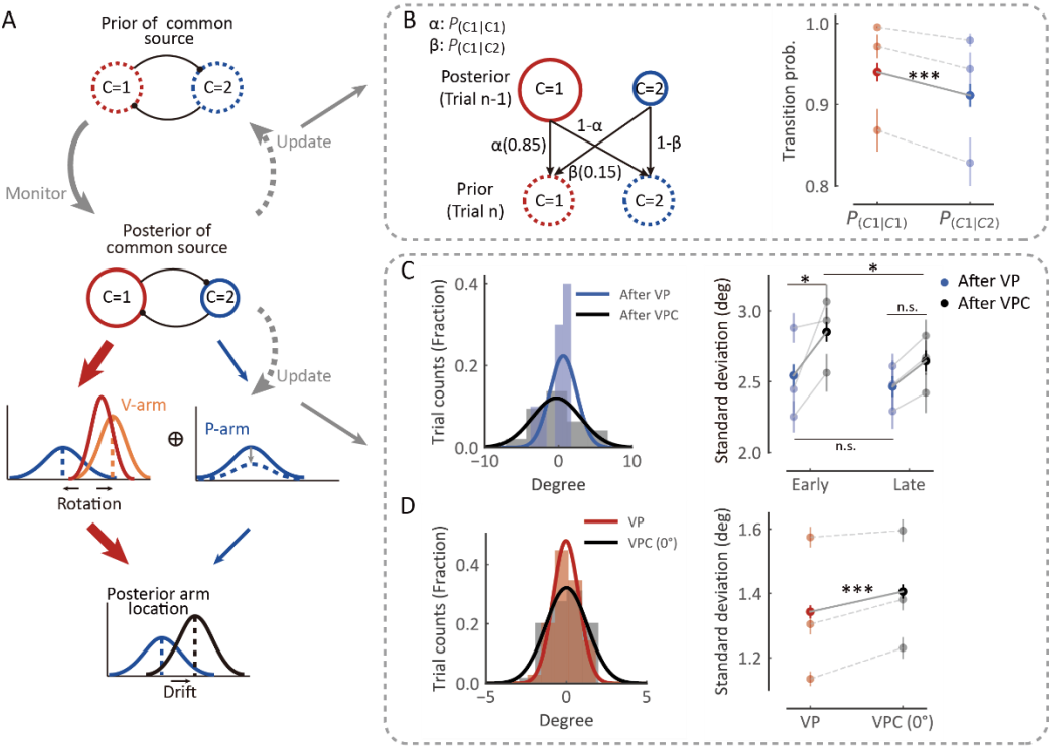
Causal inference model predicts the dynamic updating of monkey’s behavior. **(A)** Schematic drawing of the dynamic hierarchical causal inference model. V-arm, visual arm signal; P-arm, proprioceptive arm signal; C=1, both V-arm and P-arm come from one common source; C=2, V-arm and P-arm come from different sources. **(B)** Transition probability from previous trial’s *P*_*com*_ to current trial’s *P*_*prior*_. Left: the transition probability of an example session. Right: average transition probabilities across all sessions from three monkeys. The darker-colored points represent the average transition probabilities across monkeys. The lighter-colored points represent the average transition probabilities of the three monkeys separately (Wilcoxon signed-rank test, *W* = 6996.0, *p* < 0.001). **(C)** After-trial effect of sensory updating. Left: the distribution of arm locations in P blocks after VP and VPC tasks in an example session. The solid lines were fitted with Gaussian distributions. Right: the averaged standard deviations of drift in P blocks after VP and VPC tasks. The solid lines represent the averaged standard deviation of drift in VP and VPC (0°) across all sessions of all monkeys in early trials (Wilcoxon signed-rank test, *W* = 851.0, *p* = 0.012, false-discovery rate [FDR] corrected) and in late trials (Wilcoxon signed-rank test, *W* = 1,024.0, *p* = 0.073, FDR corrected). The uncertainty of P trials after the VPC task in the early part of the session was significantly larger than that in the later part (Wilcoxon signed-rank test, *W* = 917.0, *p* = 0.035, FDR corrected); this is not the case for P trials after the VP task (Wilcoxon signed-rank test, *W* = 1,086.0, *p* = 0.15, FDR corrected). **(D)** Within-trial effect of sensory updating. Left: the distribution of arm locations in VP and VPC (0°) tasks. The solid lines were fitted with Gaussian distributions. Right: the average standard deviation of drift in VPC (0°) trials was significantly higher than that in VP trials (Wilcoxon signed-rank test, *W* =10,035.0, *p* < 0.001). The dashed lines represent the average standard deviation of the drift in VP and VPC (0°) in each monkey. Error bars indicate SEMs; \**p* < 0.05; *** *p* <0.001; n.s., not significant.

We next examined whether the sensory representation is updated to maintain consistency with the causal structure of the environment. That is, the estimates of physical arm locations should tradeoff in systematic ways depending on the current common-source belief (e.g., *P*_*com*_ in different tasks: VP, P and VPC). For example, when the monkey incorrectly infers the visual and proprioceptive arms come from the same source when a disparity is presented, the uncertainty of proprioception should increase to “explain away” the conflict between the two inputs. According to this idea, since that the block design in the current experiment resulted in P trials (in the P task) sometimes following VPC task and other times following VP tasks, we then reasoned that because the overall *P*_*com*_ was lower in the VPC task than in the VP task, the uncertainty of proprioception (i.e., the distribution of proprioceptive drifts in the P trials) would be larger after the VPC task than after the VP task. We analyzed the drift variation in P trials and found that, in the early trials (first third of each P block), the uncertainty of P trials following the VPC task was significantly larger than that following the VP task (Fig. 2C, Wilcoxon signed-rank test, *p* = 0.012). The increase in the uncertainty of proprioception was recovered in the late trials (last third of each P block), evident by a significant difference in the uncertainty between early and late P trials (Fig. 2C, Wilcoxon signed-rank test, *p* = 0.035). The decrease in the uncertainty of proprioception was reasonable, as the tradeoff effect in VPC task gradually recovered. Furthermore, we hypothesized that if a tradeoff of sensory representation occurs during the process of causal inference, the tradeoff would also affect the uncertainty of VP integration in both VP and VPC tasks. We examined the distribution of proprioceptive drifts using the trials with 0° disparity in the VPC task, in which the V and P information were congruent, and compared it with the distribution in the VP task. As predicted, we found that the variance of the proprioceptive drift was significantly larger in the VPC task than in the VP task (Fig. 2D, Wilcoxon signed-rank test, *p* < 0.001). As a control, we also investigated whether the mean of drift, representing the accuracy of proprioceptive arm, was affected by the causal structure of the environment. We found there was no significant difference between the mean of drift for P trials following the VPC task and that following the VP task in both early part (Fig. S2, left, Wilcoxon signed-rank test, *p* = 0.37, FDR corrected) and late part (Fig. S2, right, Wilcoxon signed-rank test, *p* = 0.37, FDR corrected). Besides these, we also found that the mean of proprioceptive drift was not updated in the VPC task compared with VP task (Fig. S2 right, Wilcoxon signed-rank test, *p* = 0.29). Thus, these results supported the notion of a tradeoff in proprioception according to causal inference environments; that is, the uncertainty, not the accuracy, of sensory representation is updated dynamically based on the task environment (*P*_*com*_).

To summarize the above-described behavioral results, first, we found that proprioceptive drift in monkeys shows a nonlinear dependency on the disparity between proprioceptive and visual input, which was well explained by the causal inference model. Second, we showed that the *P*_*com*_ integrated with visual-proprioceptive sensory inputs and is updated by historical information in a trial-by-trial basis. Third, to maintain a consistency of causal inference, sensory uncertainty, reflected by the variance of proprioceptive drift, is updated in the inference along with the change of *P*_*com*_. Taken together, we established the behavioral paradigm in which monkeys infer the hidden cause by integrating prior information and sensory inputs while dynamically updating both *P*_*com*_ and sensory representation. The behavioral responses of the monkeys enabled us to examine the underlying neural mechanisms and functional circuits.

### Causal inference in individual premotor and parietal neurons

We recorded from two brain regions, premotor cortex (475 neurons) and parietal cortex (area 5; 238 neurons), in the three monkeys performing the reaching tasks (Fig. 3A, for details, see Supplementary Materials). We first examined the neural representations of the visual and proprioceptive arm locations in each trial during the target-holding period in the VPC, VP, and P tasks (Fig. 3B). Both brain regions conveyed significant information about the arm location in the three tasks, as measured by a bias-corrected percent explained variance (ωPEV) (Fig. 3C, Wilcoxon signed-rank test, *p* < 0.001, FDR corrected; see Methods). In the VP and P tasks in which there are no visual-proprioceptive disparities, both premotor and parietal regions showed similar visual and proprioceptive arm information (Fig. 3C, left, VP arm, Wilcoxon rank-sum test, *p* = 0.97, FDR corrected; P arm, Wilcoxon rank-sum test,, *p* = 0.49, FDR corrected).

**Figure 3.**
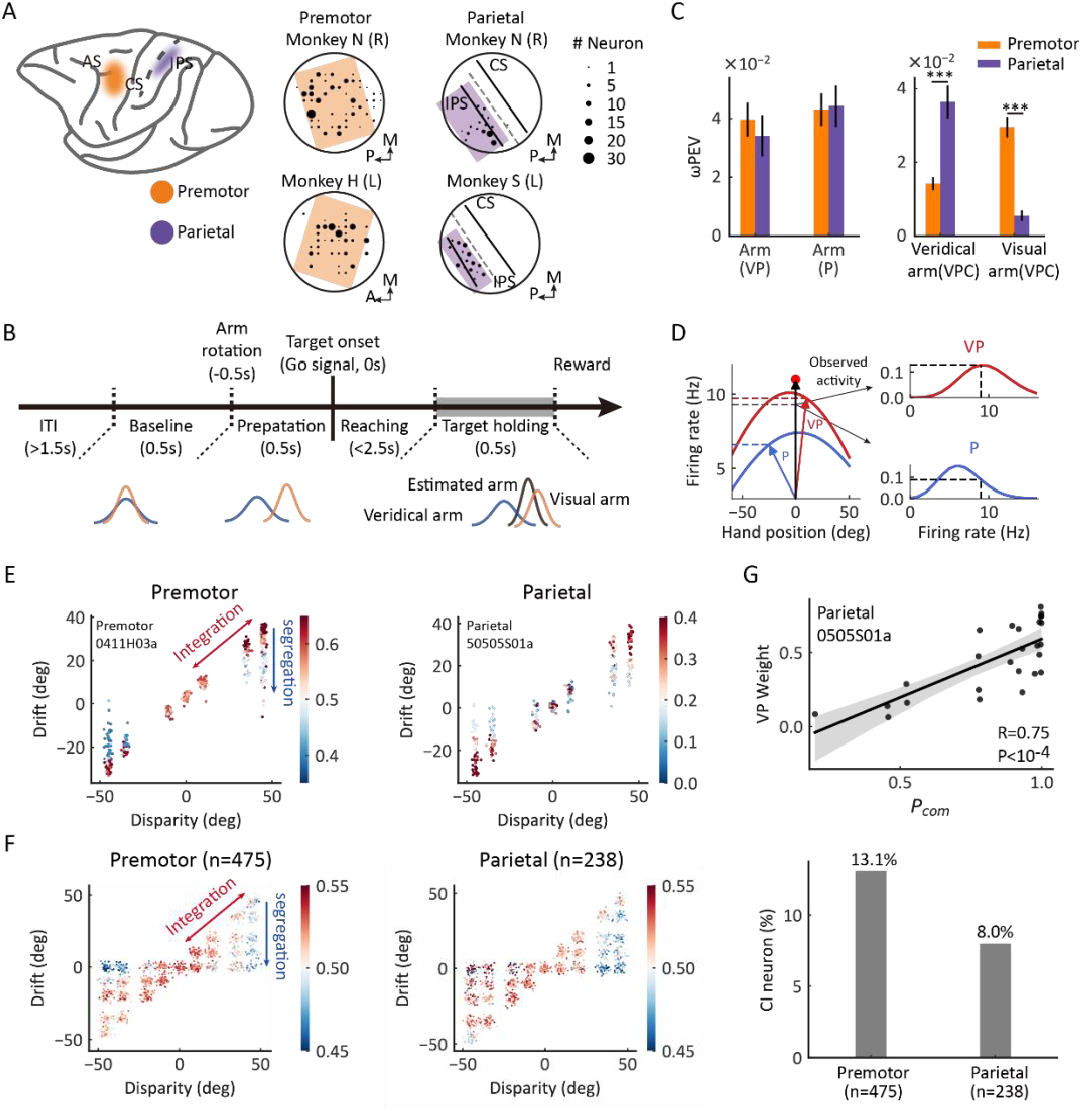
Casual inference neurons in premotor and parietal cortices. **(A)** Recording sites. Left: two regions of interest were recorded through single electrodes in macaque monkeys. Middle and right: specific recording sites in three monkeys. L, left hemisphere; R, right hemisphere; A, anterior; P, posterior; M, medial. **(B)** Temporal structure of a single trial for the VPC condition. **(C)** Neural information of arm locations in premotor and parietal cortices. Left: No significant difference between the brain regions for the neural information of VP arm (Wilcoxon rank-sum test, *W* = 0.040, *p* = 0.97, FDR corrected) and P arm (Wilcoxon rank-sum test, *W* = -0.90, *p* = 0.49, FDR corrected), respectively. Right: There were significant differences between the brain regions for both the neural information of Veridical arm (Wilcoxon rank-sum test, *W* = -5.32, p < 0.001, FDR corrected) and Visual arm (Wilcoxon rank-sum test *W* = 5.68, *p* < 0.001, FDR corrected) in VPC condition, respectively. Both brain regions conveyed significant information about the arm location in the three tasks (PMC: VP arm, Wilcoxon signed-rank test, *W* = 38,146.0, *p* < 0.001, FDR corrected; P arm, Wilcoxon signed-rank test, *W* = 34,983.0, *p* < 0.001, FDR corrected; Veridical arm (VPC), Wilcoxon signed-rank test, *W* = 35,062.0, *p* < 0.001, FDR corrected; Visual arm (VPC), Wilcoxon signed-rank test, *W* = 22,226.0, *p* < 0.001, FDR corrected. Area5: VP arm, Wilcoxon signed-rank test, *W* = 9,390.0, *p* < 0.001, FDR corrected; P arm, Wilcoxon signed-rank test, *W* = 7,324.0, *p* < 0.001, FDR corrected; Veridical arm (VPC), Wilcoxon signed-rank test, *W* = 3,552.0, *p* < 0.001, FDR corrected; Visual arm (VPC), Wilcoxon signed-rank test, *W* = 10,483.0, *p* < 0.001, FDR corrected). Error bars indicate SEMs. **(D)** Schematic drawing of VP weight analysis (see Methods) in one example trial for the VPC condition. **(E)** Two examples of causal inference neurons in premotor and parietal cortices during the target-holding period. Each point represents one single trial, and the color represents the value of VP weight. **(F)** Population causal inference patterns in two brain regions. Each point was a pseudo-trial that was generated through bootstrapping, and the color represents the value of VP weight. **(G)** An example neuron in parietal cortex showing the causal inference pattern defined by a significant positive correlation between VP weight and *P*_*com*_ (Pearson’s correlation). Each point represents the average *P*_*com*_ and VP weight in a cluster from the behavioral *P*_*com*_ pattern. The solid line was fitted with linear regression, and the shaded area indicates the 95% confidence interval. The bar plot represents the fraction of causal inference neurons in premotor (13.1%, One sample Z-test, *Z* = 5.21, *p* <0.001) and parietal (8.0%, One sample Z-test, *Z* = 1.70, *p* = 0.045) cortices which were significantly higher than chance level (5%), respectively. There was a significant difference between the brain regions (Pearson’s chi-square test, *χ*^2^ = 3.89, *p* = 0.049). *** *p* < 0.001.

However, when disparities were introduced in the VPC task, the premotor cortex showed a stronger signal for visual arm information (Fig. 3C, right, Wilcoxon rank-sum test, *p* < 0.001, FDR corrected), whereas parietal cortex showed stronger signals for information related to the proprioceptive arm (Fig. 3C, right, Wilcoxon rank-sum test, *p* < 0.001, FDR corrected). This suggests that premotor and parietal regions may play different roles during causal inference processing.

To further quantify neural information about causal inference in the VPC task at the single-neuron and single-trial levels, we utilized the VP and P tasks to characterize neural responses, as these tasks involve expected stereotypical behaviors in the two extreme regimes: full integration and segregation. Thus, neurons that are more active during the VP task reflect a preference for integrating congruent VP information and, hence, constitute a natural candidate for “integration (VP) neurons”. By contrast, neurons that are more active during the P task are likely candidates for “segregation (P) neurons”. We then implemented a linear probabilistic model which combined how the neural response pattern aligned with the VP and P response profiles and used this model to implement a probabilistic decoding analysis to calculate the probability of VP or P (VP weight = *P*_*vp*_/[*P*_*vp*_ + *P*_*p*_]) on the basis of the firing rate in each trial (Fig. 3D, also see Methods). Thus, for a single trial, a larger VP weight denotes a higher probability of integration (high *P*_*com*_). We first focused on the target-holding period in a trial, as the neurons could well display their spatial tunings when monkeys holding their arms on the target. We found that both premotor and parietal regions carry information about *P*_*com*_ at the single-neuron level during the target-holding period (Fig. 3E and F). That is, the VP weight of the neuron or population progressively decreased along with the disparity, and in trials with large disparity (e.g., 35° and 45°), the neuron(s) had a higher VP weight when the drift was large (i.e., the monkey integrated the visual information; thus, a high *P*_*com*_ predicted by the BCI model) and shifted gradually toward higher P weights when the drift shifted to 0 (i.e., the monkey segregated the visual information; thus, a low *P*_*com*_ predicted by the BCI model). The VP weight was highly correlated with the *P*_*com*_ from behavior (Fig. 3G). Note that premotor cortex had a slightly higher proportion of causal inference neurons (13.1%) than parietal cortex (8.0%, Pearson’s chi-square test, *χ*_2_ = 3.89, *p* = 0.049).

### Population states encode P_com_ during causal inference

We next focused on the overall populations of neurons in both regions and asked whether and how their population states reflect the uncertainty of causal structure, *P*_*com*_. We were guided by the results from single-neuron analyses during the target-holding period described above, in which neurons responsive to high *P*_*com*_ (prefer integration) are more likely to show neural tuning similar to that during the VP task, and neurons responsive to low *P*_*com*_ (prefer segregation) show a tuning profile similar to that in the P task. We thus hypothesized that there are neural components or subspaces embedded in the population activity that represent the dynamic change in the coding of *P*_*com*_ in the VPC task, which would lie between the components representing the VP and P profiles. Furthermore, the computation of *P*_*com*_ in the BCI model is determined by the relation and disparities between the visual information from the artificial arm and proprioceptive information from the monkey’s actual arm. In other words, according to the model, the causal inference can be constructed before the visual target appears, and the participant uses this information to guide the reach. We thus further hypothesized that the dynamics of the population states also reflect the *P*_*com*_ during the preparation period, during which there is no motor planning or preparation.

Thus, we grouped trials from each neuron into high and low *P*_*com*_ classes according to the drift under each disparity (high, top third of the trials [in red]; low, bottom third of the trials [in blue]) (Fig. 4A). We conducted demixed principal component analysis (dPCA) to visualize any neural component that represents the *P*_*com*_ in the VPC task in relation to that in the VP and P tasks (see Methods). dPCA decomposes population activity into a set of dimensions that each explain the variance of one factor of the data (Kobak et al., 2016). We included the factors of time, arm location, and *P*_*com*_ (Fig. 4B). In the analysis, VP and P trials were included, which served as the templates of integration and segregation, respectively. As shown in the schema (Fig. 4B), if the decomposed neural components indeed represent the *P*_*com*_, the population activity of high and low classes in this subspace should lie between that of the VP and P classes and the four classes (high, low, VP, and P) should be separated from each other. The dPCA results indicated that the *P*_*com*_ components, which were unrelated to the arm location, represented 29.9% and 20.5% of the total firing rate variance in the premotor and parietal areas, respectively (Fig. 4C, in red). Importantly, the activity in *P*_*com*_ dimensions seems consistent with our hypothesis, demonstrating the dynamics of *P*_*com*_ between integration (VP) and segregation (P). In addition, compared to the activity in parietal cortex, the neural trajectories of the premotor populations showed an earlier divergence in *P*_*com*_ dimensions (Fig. 4D).

**Figure 4.**
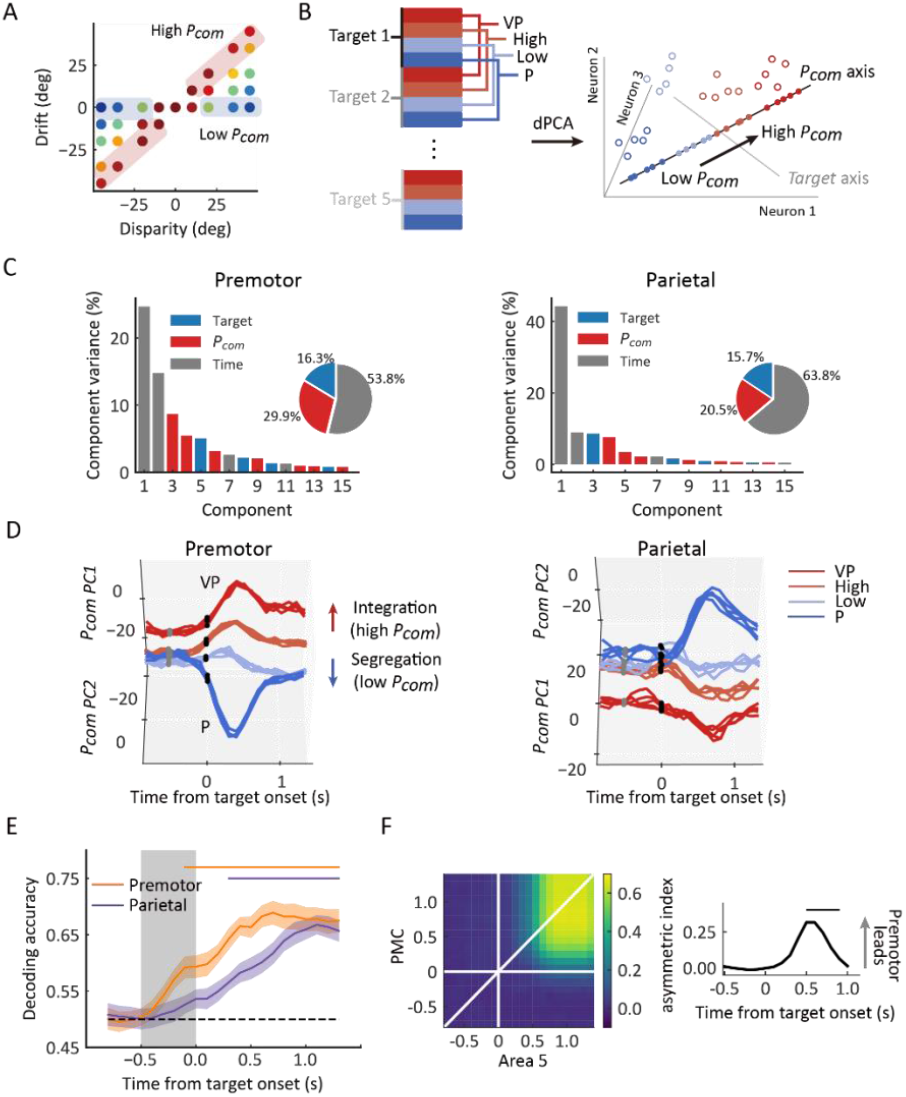
Dynamic population decoding of *P*_*com*_. **(A)** Schematic drawing of the high *P*_*com*_ group (top third of trials) and the low *P*_*com*_ group (bottom third of trials) based on the relative drift (drift/disparity). **(B)** Schematic drawing of the dPCA. All trials of each neuron were grouped into 20 classes (5 targets × 4 conditions, including VP and P tasks and high and low groups in the VPC task). The marginalization matrix was generated by averaging all trials in each class. **(C)** dPCA decomposes population activity into a set of components given the task parameters of interest. **(D)** Temporal evolution of dPCA components of *P*_*com*_. The gray points represent the disparity onset; the black points represent the target onset. **(E)** Population decoding of *P*_*com*_. The decoding accuracy was plotted as a function of time. The gray shaded area represents the preparation period. The horizontal dashed black line represents the chance level. The horizontal solid colored lines at the top represent the time of significant decoding accuracy (cluster-based permutation test, *p* < 0.05). Shaded areas indicate 95% confidence intervals. **(F)** jPECC results averaged across all sessions. Left: x-axis represents the time of Parietal (Brodmann Area 5) from target onset; y-axis: represents the time of Premotor (PMC) from target onset.The color bar represents the cross-validated correlation coefficient. Right: lead-lag interactions as a function of time relative to target onset. The horizontal black line represents the time of significant jPECC asymmetry index versus shuffled data (cluster-based permutation test, *p* < 0.05).

To further quantify their dynamics in a statistical manner, we trained a linear support vector machine (SVM) using pooled activities in each brain region through the entire trial. The dynamic decoding results showed that the *P*_*com*_ information is correctly predicted by neuronal population activities in both regions after target onset but is decoded only by premotor neurons during the preparation period, when there was no visual target or motor preparation (Fig. 4E, cluster-based permutation test, *p* < 0.05). This may suggest that the premotor cortex is where causal inference is computed and sends the information to parietal cortex during the reaching period.

Next, we tested the relationship between the population activities in the two areas. We performed a joint peri-event canonical correlation (jPECC) analysis, which detects correlations in a “communication subspace” between two brain regions(Steinmetz, Zatka-Haas, Carandini, & Harris, 2019). In brief, we conducted a canonical correlation analysis for every pair of time points containing the population neural firing rates from the two regions. If the shared neural activity emerges at different times in the two regions, that is, activity in one region potentially leads to activity in the other one, then we should observe a temporal offset between them. The jPECC results revealed a significant time lag for activity correlations between premotor and parietal areas in *P*_*com*_ dimensions (Fig. 4F, cluster-based permutation test, *p* < 0.05), suggesting a potential feedback signal of *P*_*com*_ from premotor cortex to parietal cortex. As a control, we performed the same procedure with misalignment trials (see Methods) to exclude the probability that the observed time lag resulted from the intrinsic temporal property of neuronal activities in these regions. There was no significant time lag between premotor and parietal areas when the trials were misaligned (Fig. S3).

### History-dependent P_com_ in premotor cortex

The behavioral experiments showed that the *P*_*com*_ can be updated by previous sensory experience in a trial-by-trial basis. To test the effect of the historical *P*_*com*_ on the causal inference in each trial, we examined neural activities during the baseline period in the VPC task, before a disparity in the visual and proprioceptive arm is introduced (Fig. 5A). We again classified the trials according to high and low *P*_*com*_. Figure 5A depicts the results from an example premotor neuron, showing that during the baseline period the neural activity exhibited selectivity toward the previous trial’s *P*_*com*_, and at the same time its neural trajectories in high and low prior classes lied between the VP and P templates. Of 475 neurons in premotor cortex, 39 (8.2%) showed such selectivity to the previous trial (Fig. S4).

**Figure 5.**
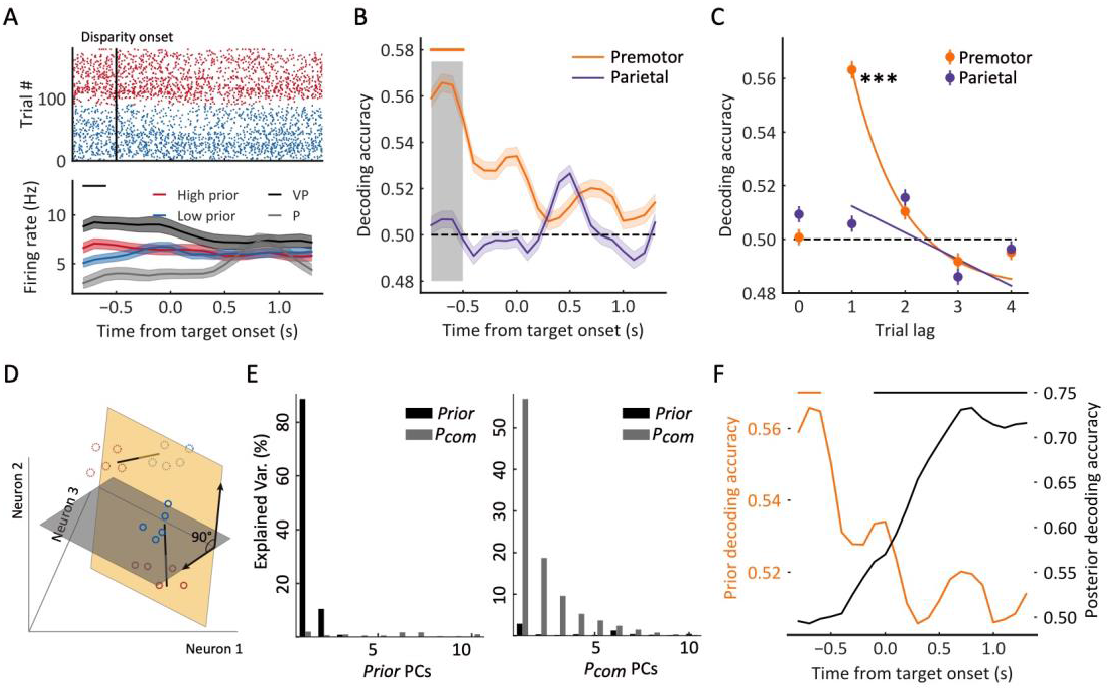
Premotor neurons encode prior information (previous trial’s *P*_*com*_) during the reference period. **(A)** Example neuron in premotor cortex showing selectivity to prior information during the baseline period. The trials in the raster plot were sorted by the *P*_*com*_ in the previous trial and grouped into high (red dots) and low (blue dots) groups. Bottom: temporal evolution of the average firing rate of “high prior” and “low prior” groups. The black horizontal line at the top represents the time window with a significant difference (two-sided *t* test, *t* = 2.36, *p* = 0.019). Shaded areas indicate SEMs. **(B)** Dynamic population decoding of prior information (n^th^ – 1 trial). The gray shaded window represents the reference period. The horizontal solid colored line at the top represents the time with significant decoding accuracy with a cluster-based permutation test (*p* < 0.05). Shaded areas indicate 95% confidence intervals. The horizontal dashed black line represents the chance level. **(C)** Decoding accuracy of prior trials (n^th^ − 1 to n^th^ − 4). Lag 0 represents the decoding of *P*_*com*_ in the current (n^th^) trial. The horizontal dashed black line represents the chance level (permutation test, *p* < 0.001). The solid lines were fitted with exponential distributions. Error bars indicate 95% confidence intervals. **(D)** Schematic drawing of orthogonal subspaces of *P*_*prior*_ and *P*_*com*_. The solid-line circles represent *P*_*com*_ and dotted circles represent *P*_*prior*_. Red represents high *P*_*com*_, blue represents low *P*_*com*_. **(E)** Left: percentage of baseline-period (*P*_*prior*_) data variance (black bars, explained variance: about 99.9%) and target-holding period data variance (gray bars, explained variance: about 10.8%) explained by the top ten prior PCs. Right: percentage of baseline-period (*P*_*prior*_) data variance (black bars, explained variance: about 13.9%) and target-holding (*P*_*com*_) period data variance (gray bars, explained variance: about 99.9%) explained by the top ten *P*_*com*_ PCs. **(F)** Premotor encoded prior information during the reference period quickly decreased after the disparity onset while the *P*_*com*_ information emerged. The orange line represents the population decoding accuracy of *P*_*prior*_ (n^th^ – 1 trial). The black line represents the population decoding accuracy of *P*_*com*_. The orange and black horizontal solid colored lines at the top represent the time with significant decoding accuracy with a cluster-based permutation test (*p* < 0.05) for prior information and *P*_*com*_ information, respectively. *** *p* < 0.001.

To further test the relation between baseline neural activity and behavior in a quantitative manner, we examined whether the population activities of these neurons can predict the *P*_*com*_ from previous trials. We trained an SVM using pooled activities across recording sessions. The historical *P*_*com*_ information was only correctly decoded from the baseline activity in premotor cortex (Fig. 5B, cluster-based permutation test, *p* < 0.05). Moreover, only recent historical information (n^th^ − 1 trial) had a significant impact on the current trial (Fig. 5C, permutation test, *p* < 0.001).

As both *P*_*prior*_ and *P*_*com*_ were represented in premotor neural activities, we wanted to examine their relationship in the neural states. We first found that there were very few neurons that responded to both information types (see Fig. S4). We then hypothesized that *P*_*prior*_ and *P*_*com*_ may be represented independently at a population level. To validate this hypothesis, we conducted PCA on the population activities during baseline and target-holding periods for *P*_*prior*_ and *P*_*com*_, respectively. If they are independent, the subspaces of *P*_*prior*_ and *P*_*com*_ will be near orthogonal, and the PCs of *P*_*prior*_ and *P*_*com*_ will capture little variance of each other(Elsayed, Lara, Kaufman, Churchland, & Cunningham, 2016). To quantify this, we projected the *P*_*prior*_ data onto the *P*_*com*_ subspace to calculate the percent variance explained by the *P*_*com*_ PCs and repeated the same procedure for the *P*_*com*_ data (Fig. 5D). The results show that the top ten *P*_*prior*_ PCs captured very little *P*_*com*_ variance, and similarly, the top ten *P*_*com*_ PCs captured very little *P*_*prior*_ variance (Fig. 5E). These results support the hypothesis that the two information types are represented independently in premotor cortex. However, such independency between *P*_*com*_ and *P*_*prior*_ could also be caused by their different temporal structures in the task. Thus, we examined their neural dynamics within a trial. Figure 5F shows the time course of decoding results of prior and posterior information, where the *P*_*prior*_ quickly decreased after the disparity onset, and at the same time, the *P*_*com*_ information increased and was retained until the end of the trial. These results demonstrated the dynamics in the computation of causal inference, where the information from the last trial is only preserved transiently and then used to integrate with sensory inputs to generate *P*_*com*_ information.

### Update sensory uncertainty of arm location in parietal cortex (area 5)

Finally, we investigated the neural activities associated with the updating of sensory uncertainty. The behavior results revealed a significantly greater uncertainty of proprioception in VP trials in the VPC task (low belief of a common source) than in the VP task (high belief of a common source) (Fig. 2C). We hypothesized that the sensory signals, which were used to make causal inference, in turn, updated their neuronal tunings to match inferred causal structure. We first examined the difference in neural tuning for arm location using the VP trials in the VP and VPC (trials with no disparity) tasks. Fig. 6A (right) shows an example neuron from the parietal cortex tuned to the center (0°) of arm location during reaching in the VP task, and the tuning range/uncertainty of the arm location was broader/lower in the VPC task. Here, for visualization purpose, we selected the time point when this neuron demonstrated the highest difference of ωPEV in the VP trials between VP and VPC tasks for the tuning calculation (Fig. 6A, left, peak delta ωPEV). The averaged dynamic spatial selectivity of all neurons revealed a significant decrease of the total spike rate variance explained by the arm location in the parietal cortex but not in the premotor cortex (Fig 6B, cluster-based permutation test, *p* < 0.05). Furthermore, at the population level, we performed the SVM decoding analysis of arm locations and found that only parietal cortex showed a significantly decreased decoding accuracy in the VPC task (Fig 6C, cluster-based permutation test, *p* < 0.05). We also confirmed that the change of decoding accurarcy in the parietal cortex was significantly larger than the change in the premotor cortex (two-way ANOVA, Condition (*VP* and *VPC 0°*) × Region (*parietal* and *premotor*), significant interaction effect, *p* < 0.05).

**Figure 6.**
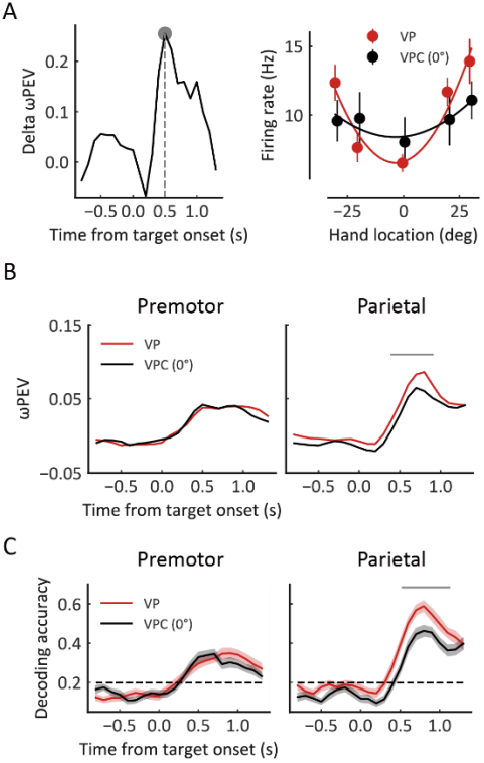
Representation of arm location is updated in parietal cortex. **(A)** Left: The difference of ωPEV between VP and VPC (0°) conditions for an example neuron in parietal cortex. Right: Snapshot of the arm location tuning for VP and VPC (0°) conditions at the time point showed in the left panel (peak delta ωPEV). The solid curves were fitted with a von Mises distribution. **(B)** Dynamic average ωPEV for VP and VPC (0°) conditions. The horizontal line at the top represents the time bins in which the ωPEV for the VPC (0°) condition was significantly lower than that for the VP condition (cluster-based permutation test, *p* < 0.05). **(C)** Dynamic population decoding of arm locations. The horizontal line at the top represents the time bins in which the decoding accuracy for the VPC (0°) condition was significantly lower than that for the VP condition (cluster-based permutation test, *p* < 0.05). Shaded areas indicate 95% confidence intervals. The horizontal dashed black lines represent the chance level.

## DISCUSSION

Our data of behavior and multi-area neural recordings revealed, for the first time, the dynamic computation of causal inference in the frontal and parietal regions at single-neuron resolution during multisensory processing. Complementary to the previous findings focused on the feedforward sequential processing of BCI, the present results demonstrate parallel top-down processing of the hidden variable of *P*_*com*_ from the premotor cortex, which monitors the weights of sensory combinations in the parietal cortex. By resolving the historical information and causal belief, the hidden causal structure and sensory representation are dynamically updated in the premotor and parietal cortices, respectively.

In the last 15 years, the BCI model has been extended to account for a large number of perceptual and sensorimotor phenomena and vast behavioral data(Shams & Beierholm). Recent studies have begun to map the algorithms and neural implementation in the human brain. Noninvasive human functional magnetic resonance imaging studies revealed a neural correlation to causal inference in parietal cortex, and magnetoencephalography showed that frontal neural activities are also involved in causal inference(Cao et al., 2019; Rohe et al., 2019; Rohe & Noppeney, 2015, HYPERLINK \l "bookmark39" 2016). However, at the single-neuron level, very few studies have examined the neural mechanism in animals. More importantly, none of the human studies have investigated the neural representation of the hidden variable, *P*_*com*_. How the prefrontal-parietal circuits contribute to the encoding and updating of *P*_*com*_ has not been explored. Our results reconciled and extended previous findings by showing that *P*_*com*_ is successively represented by premotor and parietal neural activities. Unlike previous human imaging studies, which used the final behavioral estimation as the index of causal inference(Cao et al., 2019; Rohe et al., 2019), our study directly examined the neural representation and dynamics of the hidden variable *P*_*com*_ at single-neuron and neural population levels. We showed that, even within a trial, the inference of a common source was dynamic. We thus propose a dynamic flow of information processing during causal inference, where the *P*_*com*_ is estimated from the information of sensory uncertainties and the disparity between them in the premotor cortex and then used for later sensory integration or segregation (model weighted average)(Kording et al., 2007); finally, these signals are maintained in the premotor-parietal circuit to guide the reaching behavior.

Historical experiences create our prior beliefs of the surrounding environment. It was proposed that various cognitive functions, such as sensory perception, motor control, and working memory, can be modulated by historical perception(Akrami, Kopec, Diamond, & Brody, 2018; Ernst & Banks, 2002; Rao, DeAngelis, & Snyder, 2012). Computationally, historical modulation can be well understood within the Bayesian framework(Kording & Wolpert, 2004). For instance, by imposing the BCI model in the present study, we showed that prior knowledge of a common source is updated by the hidden probability of the common source (*P*_*com*_) in the previous trial and then integrated with the sensory inputs in a Bayesian manner. The feedback from a posterior signal is one of the signatures of a hierarchical recurrent Bayesian model in a recurrent neural network(Darlington, Beck, & Lisberger, 2018). Furthermore, the posterior signal enables the construction of the causal inference environment, which can modulate sensory processing in lower-level sensory areas (e.g., parietal cortex) through a top-down feedback mechanism to maintain the belief of the causal structure. Therefore, our results provide the first behavioral and neural evidence in animals that the frontal-parietal circuit represents the hierarchical Bayesian inference and dynamically updates the causal structure and sensory representation to support the causal inference during multisensory processing.

Previous research over the past two decades has revealed that even the perceptions of body ownership and agency are remarkably malleable and involve continuous processing of multisensory information and causal inference(Kilteni, Maselli, Kording, & Slater, 2015; Legaspi & Toyoizumi, 2019). Thus, our study provides unique data toward an understanding of self-relative awareness (e.g., bodily self-consciousness) in macaque monkeys, showing neural implementation of causal inference at the neural circuit level. We also identified the hidden components of causal inference in the parietal and premotor cortices of macaque monkeys by using a visual-proprioceptive task. This is important, because, unlike most sensory cognitive functions, the subjective perceptions of body ownership and agency cannot be directly measured from explicit reports from animals. Using the BCI model and neural activities recorded from multiple brain areas, we now are able to begin exploring body ownership and agency qualitatively by examining the hidden variable in both behavior and neural representations.

In the BCI framework, there are two key components, inferring the hidden variables (e.g., *P*_*com*_) and updating the causal structure and sensory representation. We have suggested that the representation and core computation of the hidden common source most likely takes place in the premotor cortex(Ehrsson & Chancel, 2019; Fang et al., 2019), which is consistent with findings for body awareness in humans(Blanke, Slater, & Serino, 2015; Ehrsson, Spence, & Passingham, 2004). The posterior belief of a common source is calculated using a Bayesian approach by integrating prior knowledge and sensory entities, and theoretically, these components should be dynamically updated at different time hierarchies. For example, the prior configuration of the body, known as the body schema in psychology, constrains the possible distribution of the body states but is dynamically updated when the context changes to maintain consistency between the internal body model and sensory inputs (e.g., rubber hand illusion or body illusion)(Botvinick & Cohen, 1998; Kilteni et al., 2015). Pathological impairment in inferring the sensory source can result in somatoparaphrenia, in which the patient declares that his or her body part belongs to another person despite the visual and proprioceptive signals from the common source of their own body(Keromnes et al., 2019). Similarly, schizophrenia patients suffering from delusions of agency have shown impairments in updating their internal causal structures. They show a deficit in the ability to detect the source of their thoughts and actions and thus incorrectly attribute them to external agents(Haggard, 2017). Therefore, although we demonstrated the neural representations and their updating by using the multisensory and reaching task in monkeys, the computational mechanism and underlying neural circuits might contribute to learning and inference in any task that relies on causal inference.

## ACKNOWLEDGMENT

We thank Florent Meyniel and Tianming Yang for their comments on the manuscript, and Xinjian Jiang, Jian Jiang, and Juntao Feng for experimental assistance. This work was supported by the Strategic Priority Research Programs XDB32070200 and XDB32010300, the CAS Pioneer Hundreds of Talents Program, the Shanghai Municipal Science and Technology Major Project 2018SHZDZX05, and the Shanghai Key Basic Research Project 16JC14202001 to L.W.

## METHODS AND MATERIALS

### Experimental model and subject details

All animal procedures were approved by the Animal Care Committee of Center for Excellence in Brain Science and Intelligence Technology, Institute of Neuroscience, Chinese Academy of Sciences, and were described previously in detail(Fang et al., 2019). Briefly, three male adult rhesus monkeys (*Macaca mulatta*; monkey H, N, and S, weighting 6–10 kg) participated in the experiment. During the experiment, the monkeys were seated comfortably in the monkey chairs and their heads were fixed. All monkeys were implanted with chambers for recordings.

### Method details

Some of the following methods are similar to those previous published(Fang et al., 2019).

### Apparatus

The monkeys were seated in front of a chest-height table on which a lab-made virtual reality system was placed (Fang et al., 2019). During the entire experiment, the monkey’s left arm (and the right arm in the case of Monkey H, who was right handed) was placed on the system and blocked from sight. A CCD camera (MV-VEM120SC; Microvision Co., China) captured the image of the monkey’s arm reflected in a 45° mirror. This image was projected to the rear screen by a high-resolution projector (BenQ MX602, China). Therefore, when the monkey looked in the horizontal mirror suspended between the screen and the table, the visual arm image appeared to be its real arm on the table. The lower edge of the screen was aligned to the table edge. The monkey’s trunk was close to the edge of the table, and the left shoulder was aligned with the midline of the screen. By using the OpenCV graphics libraries in C++ (Visual Studio 2010; Microsoft Co., WA, USA), the arm image and the visual target were generated and manipulated. By using CinePlex Behavioral Research Systems (Plexon Inc., TX, USA), sampled at 80 Hz, the hand position was tracked and recorded. The tracking color marker was painted onto the monkey’s first segment of the middle finger, which was not visible after adjusting the light exposure settings of the video.

### Behavioral task procedures

The monkey was trained to report its proprioceptive arm location by reaching for a target in a visual-proprioceptive causal inference task (Fig. 1A) (Fang et al., 2019). The monkey initiated a trial by placing its hand on the starting point (a blue dot with a 1.5-cm diameter) for 1,000 ms and was instructed not to move. After the initiation period, the starting point disappeared and the visual arm was rotated (within one video frame, 16.7 ms) for the visual-proprioceptive conflict (VPC) condition, and the rotation was maintained for 500 ms (the preparation period). After that, the reaching target was presented as a “go” signal. The monkey had to reach the target (chosen from T1 to T5 randomly trial by trial [Fig. 1A]) within 2,500 ms and hold its hand in the target area (see as follows) for 500 ms to receive a drop of juice as the reward. Any arm movement during the target-holding period automatically terminated the trial. The rotated arm was maintained throughout the entire trial along with the arm movement. The intertrial interval (ITI) was ∼1.5–2 s, after which the monkey was allowed to start the next trial. During the ITI, the visual scene was blank. Under the VPC condition, across trials, the visual arm was randomly presented with a disparity of 0°, ±10°, ±20°, ±35°, or ±45° (+, clockwise [CW]; −, counterclockwise [CCW] direction) from the subject’s proprioceptive arm, with its shoulder as the center point. The starting point was fixed 25 cm away from the monkey’s shoulder. The target position was selected randomly trial by trial from one of five possible positions located on an arc (a ±4° jitter was added to the original position trial by trial to ensure the monkey did not perform the task by memorizing all the target positions.

Besides the VPC condition, the monkey also was instructed to perform a vision-proprioception (VP) congruent task and proprioception-only (P) task during the recording session. The only difference between the VPC and VP condition was that during the entire trial under the VP condition, the visual arm was always congruent with the proprioceptive arm. The only difference between VP and P conditions was that during the single-trial for the P condition, the visual arm information was blocked starting from the onset of the preparation period.

Each VPC block contained 55 trials in which the 9 disparities and 5 targets were randomly combined. Each VP and P block contained 27 trials in which 5 targets randomly occurred in every single trial. In one recording session, typically one or two P blocks were given first to ensure that the monkey performed the task with its proprioceptive arm, and then in the following blocks, VP, P, and VPC conditions were randomly mixed. One recording session contained more than 3 VP and P blocks and more than 8 VPC blocks.

### Target (with reward) area

To ensure the monkeys indeed performed the reaching-to-target task with their proprioceptive hand, under the VPC condition, the reaching target area (with reward) was defined as follows: the radial distance from the hand to the center of the target was less than 5 cm to ensure that the monkey did reach out to the target; with the target as the center, the azimuth range was set from [−8 + rotation degree] to +8° when the rotation degree was negative (counter-clockwise), and from –8° to [+8 + rotation degree] when the rotation degree was positive (clockwise) (green zone in Fig. 1B). Only the correct trials were used in the subsequent analysis.

### Electrophysiology

Extracellular single-unit recordings were performed described previously (Fang et al., 2019) from three hemispheres in three monkeys. Briefly, under strictly sterile conditions and general anesthesia with isoflurane, a cylindrical recording chamber (Crist Instrument Co., Inc., Maryland, USA) of 22 mm in diameter was implanted in the premotor cortex and in the parietal area 5. The location of the recording chamber on each animal was determined by individual MRI atlas (3T, Center for Excellence in Brain Science and Intelligence Technology, Institute of Neuroscience, Chinese Academy of Sciences) (Graziano, 1999; Graziano, Cooke, & Taylor, 2000; Matelli & Luppino, 2001). During the recording session, glass-coated tungsten electrodes (1–2 MΩ; Alpha Omega, Israel) were inserted into the cortex via a guide tube using multi-electrode driver (NAN electrode system; Plexon Inc., USA). On-line raw neural signals were processed offline to obtain a single unit by Offline Sorter (Plexon Inc., Dallas, TX). The sorted files were then exported to NeuroExplorer software (Plexon Inc., Dallas, TX) to generate a mat format for analysis in MATLAB (Mathworks, Natick, MA, USA) and Python (The Python Software Foundation).

### Quantification and statistical analysis

All statistical analyses were implemented with scripts written in MATLAB or Python. In premotor cortex, 475 neurons were recorded from two monkeys (272 neurons from Monkey H and 203 neurons from Monkey N); in parietal area 5, 238 neurons were recorded from two monkeys (116 neurons from Monkey N and 122 neurons from Monkey S). As all monkeys’ behavior and model fitting results were similar, for all analyses, data were combined across monkeys. All related statistics are reported in the Figure legends.

### Analysis of behavior data

#### Bayesian causal inference model

To capture the uncertainty of causal structure, the core of causal inference, the Bayesian causal inference (BCI) model described in a previous visual-proprioceptive integration study (Fang et al., 2019) was adopted. In the present study, the BCI framework included three models: (i) the full-segregation model, which assumes that visual and proprioceptive estimates of the arm’s locations are drawn independently from different sources (C=2) and processed independently; (ii) the forced-fusion model, which assumes that visual and proprioceptive estimates of the arm’s locations are drawn from a common source (C=1) and integrated optimally, weighted by their reliabilities; and (iii) the BCI model, which computes the final proprioceptive estimate by averaging the spatial estimates under full-segregation and forced-fusion assumptions weighted by the posterior probabilities of common source. Here, the BCI model assumes that both visual and proprioceptive location information (*S*_*V*_ and *S*_*P*_) are represented as *x*_*V*_ and *x*_*P*_ in the neural system, respectively, which are drawn from the normal distribution with sensory noise [*N*(*S*_*V*,_ *σ*_*V*_), *N*(*S*_*P*_, *σ*_*P*_)]. The causal inference structure is determined by the joint distribution of two sensory signals (sensory likelihood) and the prior probability of a common source (*P*_*prior*_). Thus, according to the Bayesian rule, the posterior probability of common source (one source probability [*Pcom*]) is calculated as follows:

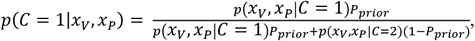

and the two sources of probability are *p*(*C* = 2 |*x*_*V*,_ *x*_*P*_) = 1 − *p*(*C* = 1|*x*_*V*,_ *x*_*P*_). If the system completely “believes” the two sensory signals are from different sources (full-segregation situation), the proprioceptive arm position is estimated independently from the visual information, as follows:

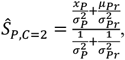

where *N*(*µ*_*Pr*_, *σ*_*Pr*_) represents a prior distribution of arm locations. In this experiment, the *µ*_*Pr*_ was set to 0 and *σ*_*Pr*_ was set to 10,000 to approximate a uniform distribution. If the system completely “believes” there is only one common source for the two sensory signals (forced-fusion situation), then the estimate of arm position is determined by the optimal integration rule, as follows:

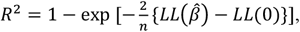

In the model simulation, the proprioceptive arm position at the end of the trial was set to zero (*S*_*P*_ = 0), so that the visual arm position is the visual-proprioceptive (*S*_*V*_ = disparity). In the task, monkeys were required to report their proprioceptive arm position, thus only the proprioceptive estimate was simulated.

#### Model fitting

To estimate the best-fitting model parameters in the BCI model, for each recording session, an optimization search was implemented that maximized the log likelihood of each model given the monkey’s data under the VPC condition. The prior probability of common source (*P*_*prior*_) and visual and proprioceptive standard deviations, σ_*V*_ and σ_*P*_, respectively, were set as free parameters to be optimized. For each optimization step, 5,000 trials per disparity were simulated to form the distribution, and the sum log likelihood of the observations given the model was calculated for each disparity. Then, the parameters were optimized by minimizing the sum log likelihood using a genetic algorithm (ga function in MATLAB). The procedure was the same as for the optimal integration model, except that there were no causal structures and only two free parameters (σ_*V*_ and σ_*P*_) need to be optimized. All simulation and optimization processes were performed in MATLAB. Only correct trials were included.

#### Model comparison

To determine the model that best explained the data at the group level using Bayesian Information Criterion (BIC), a Bayesian random-effects model comparison was used (Rigoux, Stephan, Friston, & Daunizeau, 2014). *BIC* = −2*LL* + *k × ln*(*n*), where *LL* denotes the log likelihood, *k* is the number of free parameters, *n* is the total number of data points, and *ln* is the natural logarithm. Finally, the better model was identified at the group level by the exceedance the probability based on all sessions of monkeys’ BICs (Wozny, Beierholm, & Shams, 2010).

The models’ goodness-of-fit was reported using the coefficient of determination (*R*^2^)(Fang et al., 2019),

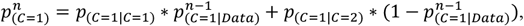

Where 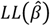 and *LL*(0) denote the log-likelihoods of the fitted and the null model, respectively, and *n* is the number of observations. The null model assumes that monkeys report the perceived arm position randomly over the disparity range from the leftmost to the rightmost. Thus, a uniform distribution over this span was predicted.

#### P_prior_ updating in causal inference

To evaluate how the historical posterior probability of common source (*P*_*com*_) influences the prior probability of common source (*P*_*prior*_), a Markov process was adopted to model the updating of (*P*_*prior*_). That is,

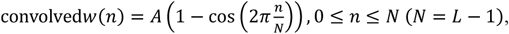

where *p*_(*C*=1)_ and *p*_(*C*=1|*Data*)_ denoted *P*_*prior*_ and *P*_*com*_ respectively, and *n* denotes the n^th^ trial under the VPC condition. Two prior states were included: C=1 (one common source) and C=2 (two different sources) at each trial. *p*_(*C*=1|*C*=1)_ denotes the transition probability from one common source (C=1) to one common source (C=1), and *p*_(*C*=1|*C*=2)_ denotes the transition probability from different sources (C=2) to one common source (C=1). For statistical significance analysis between *p*_(*C*=1|*C*=1)_ and *p*_(*C*=1|*C*=2)_, the Wilcoxon signed-rank test was used for paired data.

Note both *P*_*prior*_ and *P*_*com*_ are latent variables. During model fitting process, we first used the Bayesian causal inference model (as mentioned before) to find the overall *P*_*prior*_, *σ*_*P*_, and *σ*_*V*_ of the subjects in the day/session as the starting parameter of the subsequent Markov model. For all subsequent trials (except the first trial), both *P*_*prior*_ and *P*_*com*_ are unknown. As time goes on, starting from the first trial, the *P*_*com*_ of the current trial is obtained through the Bayesian causal inference model, and the *P*_*prior*_ of the next trial is obtained through the integration probability (*P*_*com*_) or separation probability (1 - *P*_*com*_) which are multiplied and added by the corresponding transition probability. Here, we fitted the observed data--drift to get the two free parameters transition probability. Through the transition probability, we define the influence of the *P*_*com*_ of the previous trial on the *P*_*prior*_ of the next trial.

#### Updating of proprioceptive representation

To evaluate whether the primary sensory representation was modulated by the belief of causal structure, the proprioceptive variance within and after VPC tasks was compared to the baseline condition. For the within effect, the proprioceptive drift was calculated using the trials with 0° disparity in the VPC task and trials in VP task (baseline condition). Here, the standard deviation (SD) of proprioceptive drift was used as a measurement for the reliability of proprioceptive representation, in which higher SD indicates lower reliability and *vice versa*. The mean of the proprioceptive drift for each target was normalized to zero. For the after effect, the SDs of proprioceptive drift under the P condition were compared between after the VP condition (baseline condition) and after the VPC condition. To characterize the temporal dynamic of the proprioceptive updating (after effect), trials in the first third and in the last third of the P task were compared. As control, similar analysis was conducted for the raw mean of proprioceptive drift (Fig S2). For statistical significance analysis, Wilcoxon signed-rank test was used for paired data.

### Preprocessing of single-unit data

To estimate continuous time-dependent firing rates, timestamps of spiking events were resampled at 1 kHz and converted into binary spikes for single trials. Spike trains were then convolved with a symmetric Hann kernel (MATLAB, MathWorks),

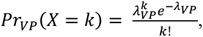

where *A* is a normalization factor ensuring the sum of the kernel values equals 1. Window width *L* was set to 300 ms. Single neurons were included in the analysis only if they had been recorded for a full set of conditions (VP, P, and VPC conditions with 9 disparities: 0°, ±10°, ±20°, ±35°, and ±45°).

Peri-stimulus time histograms (PSTHs) were then calculated for four epochs of interest in a trial: (i) the baseline epoch (500 ms before the onset of visual arm rotation), the preparation epoch (500 ms after the onset of the visual arm rotation), (iii) the target onset epoch (1,000 ms after the onset of target onset), and (iv) the target-holding epoch (500 ms after the onset of target holding). To smooth the firing rate at each time point, the neural firing rate was calculated by averaging in sliding windows (window size, 400 ms; step size, 100 ms) in a single trial, resulting in 22 time bins of mean firing rate for every single trial for subsequent dynamic analysis.

### Causal inference neuron

To measure the representation of a single neuron for causal inference on a single trial, the probability that a single neuron would integrate or segregate the sensory information on a single trial was calculated(Fang et al., 2019). The basic assumption here is that in a single trial under the VPC condition, if the neuron is more inclined to represent integrated information, then its firing rate will be closer to its response under VP conditions and the farther away from the response under P conditions, and *vice versa*. The normalized weight of integration (VP weight) was calculated as follows:

1. First, obtain the neuron response to the arm position under P and VP tasks and fit the von Mises distribution to get the tuning curve.
2. Under VPC conditions, obtain the current visual arm and the real arm positions, and at the same time, obtain the neuron’s firing rate when the arm is in the corresponding position under VP and P conditions, *λ*_*VP*_ and *λ*_*P*_, respectively.
3. The VP and P templates can be generated through the Poisson distribution:

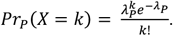
4. According to the corresponding probabilities, *Pr*_*VP*_ and *Pr*_*P*_ in the two templates are obtained, and the integration weights for this neuron in the VPC task can be obtained through standardization:

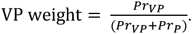

To quantitatively describe whether a single neuron is encoding causal inference, the correlation between *P*_*com*_ and VP weight is calculated. The logic is as follows: the *P*_*com*_ can be used to measure the degree of integration of sensory information and separation of sensory information at the behavioral level, whereas VP weight can measure this characteristic at the electrophysiological level. Therefore, if a neuron is performing causal inference, there should be a significant positive correlation between the *P*_*com*_ and VP weight for the corresponding behavior. Neurons that (i) respond to VP/P conditions and (ii) for which *P*_*com*_ and VP weight are significantly positively correlated in the final holding stage are called causal inference neurons. The specific algorithm was as follows:

1. First, obtain neurons with significant selectivity under VP and P conditions (ANOVA, main effect, *p* < 0.05).
2. According proprioception drift, all trials were divided into 29 classes. Continuous drift values were grouped into nine clusters: < -35°, [-35° -25°], [-25° -15°], [-15° -6°], [-6° +6°], [+6° +15°], [+15° +25°], [+25° +35°], > +35°. To be noticed, ±6° covers approximately 99% of drift distribution under the VP and P condition. Thus, for the disparity 0°, there was only one cluster [-6° +6°]. Since the distribution of drift becomes wider (higher variance) along with the larger the disparity, the more clusters would be assigned for big disparity. For example, for the disparity ±45°, there were five clusters of drifts. *P*_*com*_ and VP weight were assigned for each class by averaging all trials within it. the Pearson correlation coefficient was then calculated between *P*_*com*_ and VP weight. If the *P*_*com*_ and VP weight were correlated significantly and positively (*p* < 0.05 and *r* > 0), the neuron was called as a causal inference neuron.

To evaluate whether the fraction of causal inference neurons was significant highly than chance level (5%), one sample Z-tests (one-sided) were conducted for each brain region, respectively.

### Population pattern of causal inference

To visualize the VP weight pattern at the brain region level, the VP weight of each trial of a single neuron under VPC conditions was calculated and then divided into 29 clusters as described above. Then, the bootstrap method was used to randomly select 50 trials from each cluster for averaging. This was repeated 50 times to obtain the VP weight (50 × 29) of a neuron for visualization. This results in a 50 × 29 × *N* matrix, where *N* indicates the number of neurons in each brain region. The trial corresponding to each neuron was averaged to obtain a 50 × 29 matrix. The VP weights of a brain region were visualized in a heatmap.

### High/low P _com_ groups

To characterize the dynamic representation of the *P*_*com*_ in the entire session, all trials in a recording session were divided into high *P*_*com*_ trials and low *P*_*com*_ trials on the basis of the relative proprioception drift (RD). The relative proprioception drift (RD = drift/disparity) of each trial was calculated. The basic idea was that the larger the *P*_*com*_, the more likely the monkey was to integrate the visual and proprioceptive information, and the corresponding RD is closer to 1. The top third and bottom third of the trials were designated the high *P*_*com*_ class and the low *P*_*com*_ class, respectively. These grouping methods were verified by the demixed principal component analysis (dPCA).

### dPCA

The method for dPCA was adopted from that published in a previous study (Kobak et al., 2016). Time, target position/arm location (−30°, −20°, 0°, 20°, and 30°), and *P*_*com*_ (VP, P, high *P*_*com*_ and low *P*_*com*_) were combined to obtain the marginalized covariance matrix of the three. The neurons whose trial number was not less than 5 under a single condition were selected for dPCA. Population activity was then projected on the decoding axes and ordered by their explained total variance for each marginalization.

### Information encoded by individual neurons

The percentage of explained variance (PEV) (Buschman, Siegel, Roy, & Miller, 2011) was used to measure the basic task components encoded by a single neuron, in which PEV reflected the degree to which the variance of a single neuron can be explained for a specific task component. Generally, PEV can be expressed as a statistical value of ***η***^2^, that is, the ratio of the variance between groups to the total variance. As the statistical value of ***η***^2^ has a strong positive bias for a small sample, the unbiased ***ω***^2^ statistical value (ωPEV) (Olejnik & Algina, 2003) was used.

To evaluate the information about the locations of the veridical arm, visual arm, and estimated arm encoded by a single neuron in the VPC task, an analysis of covariance was used to decompose the variance, and the ωPEV was calculated. In detail, for a single neuron, ωPEV was calculated for each type of arm when setting other two types of arm locations as covariates. the whole reaching space was divided into 11 parts from, −45° to 45°, to transform it from a continuous variable to a discrete variable. For statistical significance analysis comparing two brain regions, a nonparametric Wilcoxon rank-sum test was used for unpaired data.

The ωPEV was calculated in each time bin to characterize the temporal dynamics of ωPEV under VP and VPC (0°) conditions. The baseline was defined as the period 500 ms before the onset of visual arm rotation, and the time bins significantly different from the baseline were determined by a one-sided, paired Wilcoxon signed-rank with false-discovery rate (FDR) correction. The time bins showing significant differences between VP and VPC (0°) conditions were determined by a cluster-based permutation test(Gramfort et al., 2013).

### Population decoding analysis

#### Decoding of P _com_

The population decoding analysis of *P*_*com*_ was performed by the linear support vector machine (SVM) classifiers with the scikit-learn toolbox(Pedregosa et al., 2011). All neurons were included in this analysis without considering their *P*_*com*_ selectivity. The classifier was trained to classify the *P*_*com*_ (high/low *P*_*com*_) with neural activity (peri-stimulus time histograms) from each brain region. All recording sessions were pooled to form a pseudo-population. Neurons with more than 50 trials in each *P*_*com*_ group were included in this analysis. Tenfold cross-validation was then implemented by splitting the neural data into 10 subsamples, each randomly drawn from the entire dataset. Decoders were then trained on 9 of the subsamples and tested on the remaining one. This process was repeated 10 times to obtain the decoding accuracy by averaging across all 10 decoders. This cross-validation process was repeated 1,000 times, and the overall decoding accuracy was taken as the mean across the 1,000 repetitions. The decoding analysis was conducted for all time points. The significance for decoding accuracy was determined by comparing the mean decoding accuracy to the null distribution from the shuffled data. The significant time duration was determined using a cluster-based permutation test for multiple comparisons across time intervals(Gramfort et al., 2013).

To test whether premotor cortex neurons encode *Pcom* earlier than area 5, a randomization test was performed between them. The corresponding numbers (here, 50 neurons per region) of neurons were randomly exchanged between the paired regions 1,000 times to generate a null distribution (chance level) of time lags, and the significance was determined by a permutation test of the true time lag from the original data and the null distribution.

#### Decoding of P_prior_

Neurons with more than 50 trials in each *P*_*com*_ group (high and low *P*_*com*_ groups, same as for the *P*_*com*_ decoding analysis described above) were selected or the *P*_*prior*_ updating decoding. The decoding procedure was the same as described for “*Decoding of P*_*com*_” unless the trials were sorted and labeled by the previous trial’s *P*_*com*_ (n^th^ trial 1 to n^th^ trial − 4) under the VPC condition. The statistical significance was determined by a cluster-based permutation test(Gramfort et al., 2013).

#### Subspace overlap analysis

PCA was performed on neural activities during the baseline period and during the target-holding period. The first ten PCs during each period were used to obtained the *P*_*prior*_ and *P*_*com*_ subspaces. To test the overlap of these subspaces, the baseline-period activity was projected onto the *P*_*prior*_ subspace, and the percent variance explained relative to the total variance of the baseline period data was quantified; similarly, the target-holding period activity was projected onto the *P*_*com*_ subspace, and the percent variance explained relative to the total variance of the target-holding period data was quantified(Elsayed et al., 2016).

#### Decoding of arm locations

All arm locations were separated into 5 spatial bins: −30°, −20°, 0°, 20°, and 30°. The basic decoding procedure was the same as described above for “*Decoding of P*_*com*_.” Neurons with more than 6 trials in each arm location bin were selected. Leave-one-out cross-validation was then implemented, and this process was repeated 1,000 times to obtain the averaged decoding accuracy. The decoding analysis was conducted for all time points. Statistical significance for decoding accuracy was determined by comparing the mean decoding accuracy to the null distribution from shuffled data. The time bins with significant difference between conditions (VP and VPC [0°]) were determined by the cluster-based permutation test for multiple comparisons across time intervals(Gramfort et al., 2013).

### Joint peri-event canonical correlation (jPECC) analysis

To test the relationship between population activities in the two brain regions, the jPECC method described in a previous study (Steinmetz et al., 2019) was utilized. First, the neuronal responses in two brain regions under the same behavior conditions, namely, high *P*_*com*_ and low *P*_*com*_, were aligned. Then, a PCA was conducted across time and trials to reduce the dimensionality to obtain the first 10 principal components (PCs) for each brain region. The trials were then divided into ten equal parts (training set and testing set) for cross-validation (10-fold cross-validation). The PCs of the training set of each brain region were used to perform a canonical correlation analysis to obtain the first pair of canonical correlation components (L2 regularization, λ = 0.5). Then, the PCs of the testing set from each brain region were projected onto the first pair of canonical correlation components, and the correlation was determined by the Pearson correlation coefficient between these projections from each region. This analysis was performed for each pair of time bins to construct a cross-validated correlation coefficient matrix. Fifty trials for each group (high *P*_*com*_ and low *P*_*com*_) from each brain region were randomly selected by bootstrapping in this analysis. Finally, a heatmap was obtained by averaging the correlation coefficient matrix repeated 1,000 times.

To quantify the lead–lag relationship of information exchange between brain regions, an asymmetric index was calculated by diagonally slicing the jPECC matrix from +300 ms to +300 ms relative to each time point (Steinmetz et al., 2019). For time point *t*, the average correlation coefficient across the left half of this slice (that is, the average along a vector from [*t* − 300, *t* + 300] to [*t, t*]) was subtracted from the right half of this slice (from [*t, t*] to [*t* + 300, *t* − 300]) to yield the asymmetry index. To test the leading significant time point across brain regions, the data from neurons in these brain regions were exchanged, and the above-described analysis was repeated 1,000 times to obtain the null distribution of the asymmetric index. Then, a cluster-based permutation test was performed to test whether the symmetric index was significantly greater than the chance level(Gramfort et al., 2013).

To further exclude the possibility that the observed lead–lag relationship resulted from the intrinsic properties of neuronal activities rather than the encoded information in these regions, all trials in each brain region were shuffled to ensure that the inter-region trials were not aligned. Then, the analysis was repeated as described above to obtain the asymmetric index.

## DATA AND CODE AVAILABILITY

Raw electrophysiology recording files, due to their size (multiple terabytes), are available upon reasonable request.

## SUPPLEMENTARY INFORMATION

**Table S1.** Model parameters and fitting evaluations of two models for monkeys.

| Subject | Causal inference (model averaging) |  |  |  |  |  | Forced fusion |  |  |  |  |
| --- | --- | --- | --- | --- | --- | --- | --- | --- | --- | --- | --- |
| | relBIC<sub>Group</sub> | EP | R<sup>2</sup> | $\sigma_P$ | $\sigma_V$ | $P_{prior}$ | relBIC<sub>Group</sub> | EP | R<sup>2</sup> | $\sigma_P$ | $\sigma_V$ |
| Monkey H | 0 | >0.999 | 0.96±0.0017 | 7.72±0.14 | 5.83±0.090 | 0.999±0.0004 | 329.52 | 2.31E-14 | 0.93±0.0039 | 9.87±0.24 | 9.02±0.23 |
| Monkey N | 0 | >0.999 | 0.93±0.0080 | 9.56±0.16 | 4.93±0.15 | 0.86±0.026 | 812.34 | 4.04E-28 | 0.37±0.36 | 11.34±0.10 | 10.46±0.13 |
| Monkey S | 0 | >0.999 | 0.96±0.0022 | 8.98±0.14 | 5.72±0.22 | 0.98±0.012 | 290.04 | 7.57E-23 | 0.94±0.0027 | 10.10±0.14 | 8.34±0.17 |

The model parameters and $R^2$ were averaged across days for monkeys; data are presented as the means ± the standard errors of the means. The relBIC<sub>group</sub> was the summation of all days' BIC for monkeys.
Abbreviations: $\sigma_P$ , standard deviation of the proprioception likelihood; $\sigma_V$ , standard deviation of the vision likelihood; $P_{prior}$ , prior probability of common source; relBIC<sub>group</sub>, Bayesian information criterion at the group level; EP, exceedance probability; $R^2$ , coefficient of determination.

**Figure S1.**
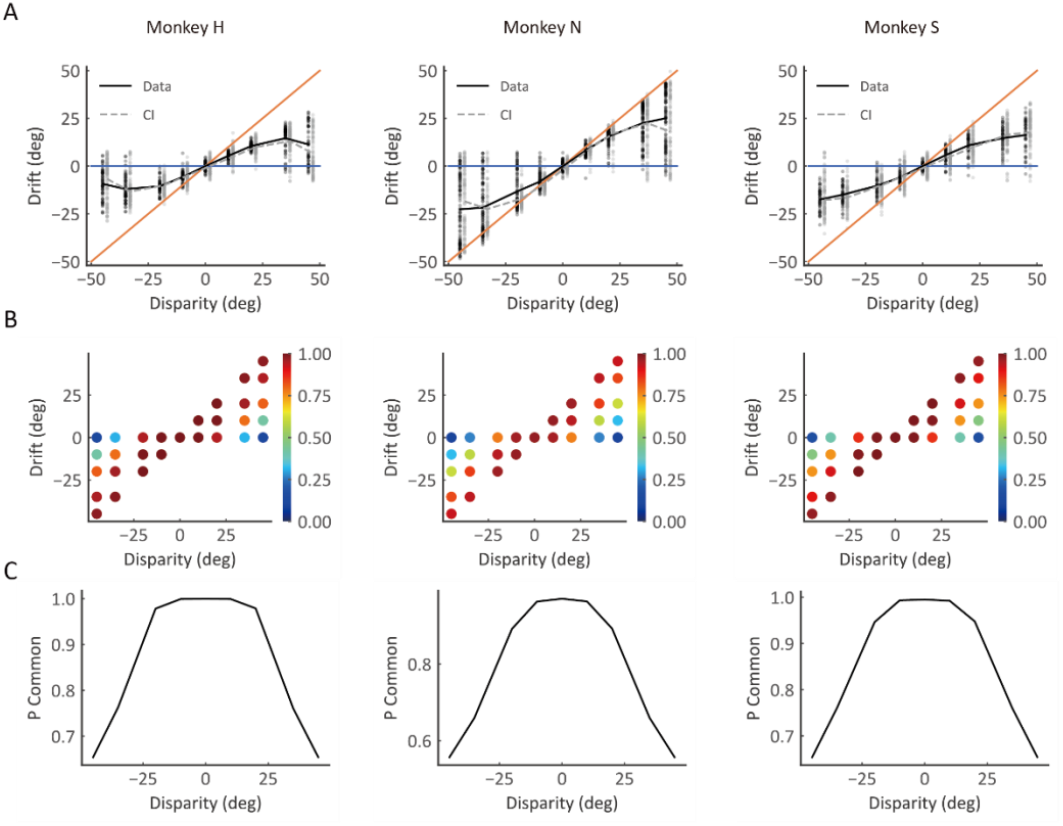
Behavior performance and causal inference model predict results in individual monkeys. **(A)** The pattern of drift is consistent across all three monkeys (black lines and dots), and the predictions of the causal inference model (gray lines and dots) characterized monkeys’ behavior data. Each dot represents a single trial, and lines represent the average result. The blue and orange solid lines represent the visual and proprioceptive bias, respectively. **(B)** Model prediction of the posterior probability of common source (*P*_*com*_). Each dot represents the averaged *P*_*com*_ in a cluster grouped by the disparity and drift based on the monkey’s behavior. **(C)** Average *P*_*com*_ as the function of disparity. The black lines represent the average *P*_*com*_ of each monkey.

**Figure S2.**
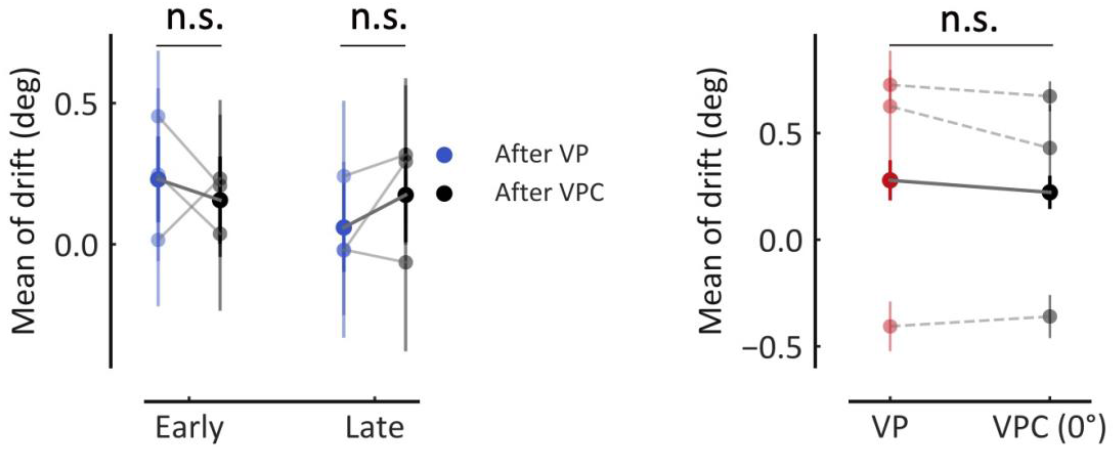
Sensory updating is not reflected in the mean of drift. Left: mean drift in P blocks after VP and VPC tasks. The solid lines represent the means of drift in VP and VPC (0°) tasks across all sessions of all monkeys in the early part of the sessions (Wilcoxon signed-rank test, *W* = 3,591.0, *p* = 0.37, FDR) and the late part of the sessions (Wilcoxon signed-rank test, *W* = 3749.0, *p* = 0.37, FDR). Right: mean of drift in VPC (0°) trials was not significantly different from that in VP trials (Wilcoxon signed-rank test, *W* = 13,668.0, *p* = 0.29). The dashed lines represent the means of the drift for VP and VPC (0°) tasks in each monkey. Error bars indicate the SEMs. n.s., not significant.

**Figure S3.**
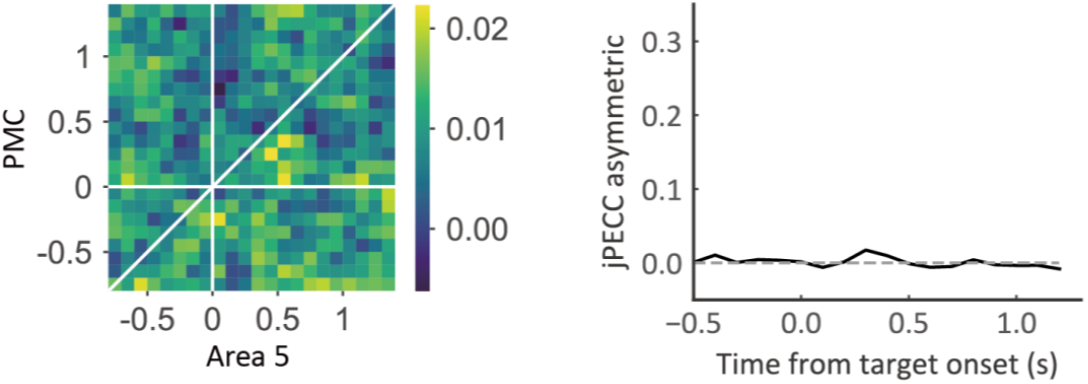
jPECC analysis with shuffled temporal alignment trials. For determining whether correlations occur with a temporal offset between premotor and parietal cortices after shuffling the trials’ alignment. Left: cross-validated correlation coefficient between premotor and parietal cortices. The trial’s temporal alignment was shuffled to determine whether correlations occur with a temporal offset between the paired brain regions. Right: the black line represents the lead–lag interactions as a function of time relative to target onset, and the gray dashed line represents the chance level (chance level = 0).

**Figure S4.**
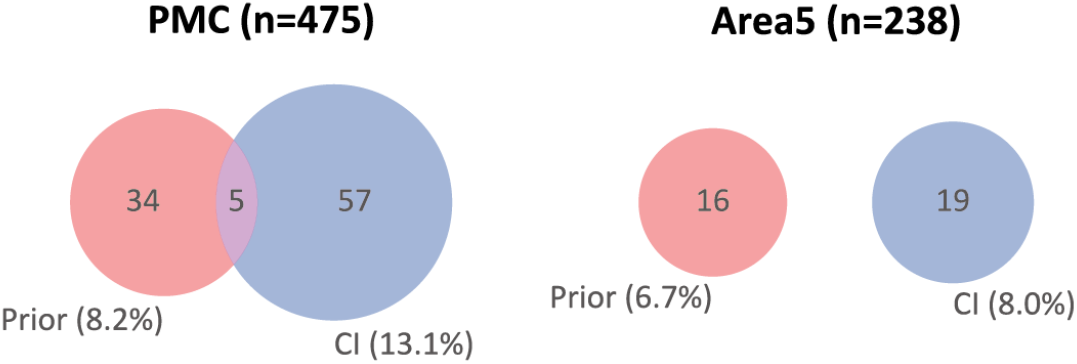
Percentage of prior-selective neurons and causal inference (CI) neurons. Red, the number of pure prior-selective neurons; blue, the number of pure CI neurons; purple, the number of dual-selective neurons.

